# Engineering dynamic interkingdom communication between synthetic organelles and mammalian systems

**DOI:** 10.64898/2026.11.25.769173

**Authors:** Satyajit Hari Kulkarni, Tiffany Rennells, Hannah Jabiro, Ashley V. Makela, Ainsley Tandoc, Austin M. Huckins, Daisuke Matsumoto, Roksana Riddle, Jonathan W. Hardy, Christopher H. Contag

**Affiliations:** Institute for Quantitative Health Science and Engineering, Michigan State University, East Lansing, MI; Department of Comparative Medicine and Integrative Biology, Michigan State University, East Lansing, MI; College of Veterinary Medicine, Michigan State University, East Lansing, MI; Department of Biomedical Engineering, Michigan State University, East Lansing, MI; Okemos High School, Okemos, MI; Department of Microbiology, Genetics, and Immunology, Michigan State University, East Lansing, MI

**Author notes:** University of Texas MD Anderson Cancer Center, Houston, TX. University of Michigan, Ann Arbor, MI. University of Pittsburgh, Pittsburgh, PA. Tokyo Metropolitan InsCtute of Medical Science, Tokyo, Japan (Daisuke Matsumoto).

## Abstract

Engineering intracellular microbes to function as synthetic organelles, or quasiorganelles, opens new avenues for controlling mammalian cell fates and function through programmable and quantitative interkingdom communication. Here, we develop a bioluminescent reporter platform to quantify the magnitude, kinetics, persistence, and spatial distribution of transcriptional regulation in mammalian host cells directed by quasiorganelles. Using an attenuated, flavin-auxotrophic *Listeria monocytogenes* chassis (LINQ1) engineered to deliver synthetic transcription factors, we demonstrate approximately 100-fold activation of mammalian reporter genes, with distinct kinetics and dynamics for the tetracycline transactivator and Gal4 systems. In transgenic reporter mice, LINQ1 induced approximately 25-fold reporter activation across multiple organs, with activity persisting beyond three weeks in some animals. In mice bearing triple-negative breast tumors, LINQ1 activated intratumoral transcription and, incidentally, transiently suppressed tumor growth, although responses diminished with repeated administration. These findings establish a quantitative foundation for engineering interkingdom communication in vivo to advance a new paradigm in which programmable microbial systems function as synthetic organelles to direct mammalian cell fates and function for therapeutic benefit.

## Introduction

Engineering communication between organisms from different kingdoms of life offers new opportunities to control cellular behavior and develop biological systems for therapeutic applications. In nature, endosymbiotic relationships have profoundly shaped the evolution of complex life, giving rise to organelles that integrate specialized functions into eukaryotic cells. Inspired by these relationships, synthetic biology offers the possibility of engineering intracellular microbes, endosymbionts, as programmable synthetic organelles, or quasiorganelles, that communicate directly with mammalian host cells to control genetic circuits.^1,2^ These engineered endosymbionts can be designed to produce and deliver transcription factors and other proteins that regulate signaling pathways, establishing a means of directing mammalian cell functions without modifying the host genome.^1,2^ Unlike conventional genetic engineering approaches, quasiorganelles offer the potential for dynamic, tunable, and reversible regulation of multiple cellular pathways through genetically encoded programs operating within the host cell. Rational engineering of synthetic organelles requires a quantitative framework for defining where, when, to what extent, and for how long these independently programmable biological systems regulate mammalian cellular functions.

Reporter systems have transformed our ability to investigate biological processes by enabling quantitative, longitudinal measurements of gene expression, cellular signaling, and molecular interactions in living systems.^3,4^ In particular, fluorescent and bioluminescent reporters have been valuable for revealing the spatial and temporal dynamics of biological activity, from individual cells to intact organisms.^3,5^ Specifically in engineered microbial systems, reporters have been useful in monitoring bacterial localization, viability, proliferation, and activation of genetic circuits in response to environmental signals.^6,7^ By coupling molecular signals produced by synthetic organelles to reporter gene activation within mammalian host cells, synthetic transcriptional circuits could provide a quantitative framework for defining the efficiency, dynamics, and spatial organization of engineered interkingdom communication.

A potential application of programmable quasiorganelles is in cancer therapy, where tumors evolve resistance to single-agent therapies through changes in tumor cell behavior and the tumor microenvironment (TME). To date, *Bacillus Calmette-Guérin* (BCG) is the only FDA-approved bacterial-based cancer therapy, and it is currently the gold standard for non-muscle invasive bladder cancer.^8^ Building on the therapeutic potential of bacteria in cancer treatment, programmable quasiorganelles could drive long-term changes in the TME^9,10^ through their intrinsic ability to localize and persist in tumors^11–13^ and activate both innate and adaptive immune responses. Given the shared nature of the TME across cancers that originate from many organ sites, such modulators could have a broad impact on cancer treatment. Additionally, the genetic programmability of prokaryotes and their capacity to express large, complex genetic payloads creates an opportunity to activate multiple pathways simultaneously to achieve durable tumor control. In the TME, macrophages are largely identified to be in a pro-tumor phenotype, producing cytokines (e.g., IL-10, TGF-β), chemokines (e.g., CCL18, CXCL8), growth factors (e.g., VEGF, FGF), and matrix metalloproteinases to drive immunosuppression, angiogenesis, stromal remodeling, and tumor growth and metastasis.^14,15^ The heterogeneity of macrophage cell surface receptor expression and phenotypes complicates therapeutic targeting by conventional approaches like antibody-drug conjugates.

We have previously developed engineered endosymbionts (EES) that target phagocytic immune cells, including tumor-associated macrophages (TAMs),^1,2^ which are a critical subset of cells in the TME and contribute to therapeutic failure.^14,16–20^ Using *Bacillus subtilis* as an EES chassis, we demonstrated the delivery of engineered mammalian transcription factors (TFs) to TAMs and subsequent modulation of mammalian cell phenotypes.^1,2^ In vitro, these endosymbionts reprogrammed macrophages towards a pro-inflammatory phenotype. In an aggressive mouse model of triple-negative breast cancer, EES treatment significantly suppressed tumor growth compared to controls.^2^ However, additional refinements to the host-quasiorganelle interaction could be made to improve cell fate reprogramming in cells and living tissues. Since *B. subtilis* produces many extracellular proteases, reducing the effective secretion of functional proteins,^21^ and is cleared by autophagy in macrophages,^2^ we created a second iteration of quasiorganelle built on an obligate intracellular strain of *Listeria monocytogenes* (Lm),^22,23^ termed ***L****isteria* **in**tracellular **q**uasiorganelle version-1 (LINQ1).

Lm is a Gram-positive, non-spore forming, motile, facultative intracellular bacterium. Lm can enter both phagocytic and non-phagocytic cells, escape the endosome by expression of listeriolysin O (LLO), and then evade the immune system by cell-to-cell spread.^24,25^ Genetic tools to engineer Lm have been extensively characterized, which allows rapid generation of recombinant strains.^26^ Additionally, Lm has been developed extensively as an anti-cancer immunotherapy in mouse models, dogs, and humans.^27–35^ Unlike *B. subtilis*, LINQ1 is capable of persisting inside macrophages by avoiding autophagy^36^ and does not significantly degrade secreted heterologous proteins, indicating lower protease production. In this study, we developed a synthetic transcriptional reporter system for engineered interkingdom communication and revealed the dynamics of transcriptional modulation by quasiorganelles from the cytoplasm of host cells (Fig. 1A,B). Using the tetracycline transcriptional activator (tTA) and Gal4 synthetic transcription factors,^37,38^ we demonstrated a maximal activation of ∼100-fold of a reporter gene in multiple cell lines and measured the dynamics across chassis organisms. In a transgenic reporter mouse, we observed significant activation compared to controls, with peak activation occurring 24-hours (h) post-treatment localized to the proximal intestinal tract and lungs. We showed that reporter mice with subcutaneous breast tumors produced similar dynamics and revealed potential therapeutic resistance following repeat treatments. This system creates a foundation to develop synthetic organelles and engineered interkingdom communication to control the fates and function of mammalian cells.

**Figure 1.**
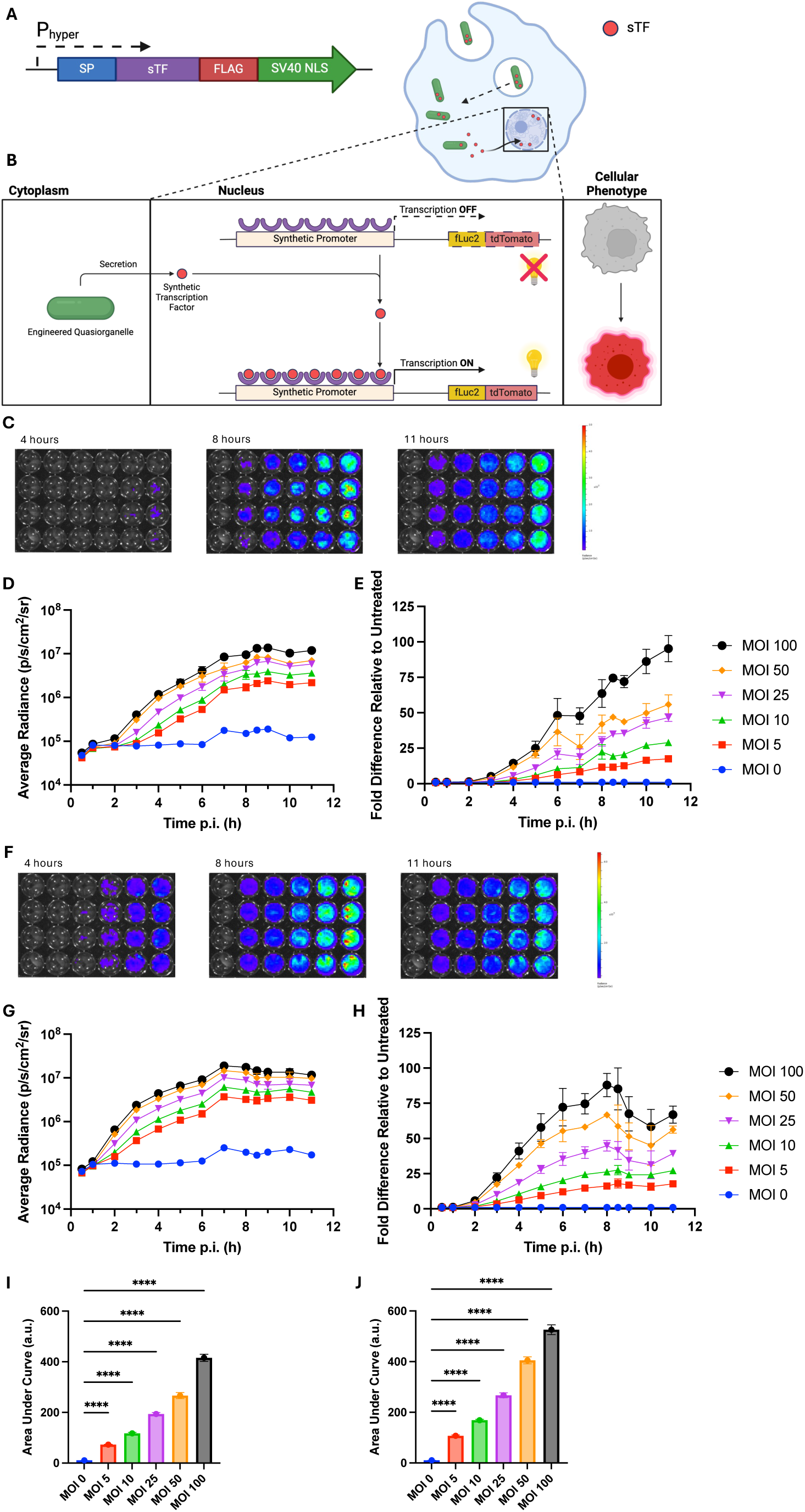
A synthetic transcriptional reporter system reveals the kinetics and dynamics of interkingdom communication between synthetic organelles and mammalian host cells. (A) Schematic of the construct expressed by the synthetic organelle, where the transcription factor has a constitutive promoter (pHyper), signal peptide (SP), the sTF gene, a 3x FLAG tag for protein detection, and an SV40 nuclear localization sequence (NLS). (B) Schematic of the reporter system, where the synthetic organelle produces a synthetic transcription factor within the cytoplasm of the host cell, which localizes to the nucleus and activates transcription of a reporter firefly luciferase (fLuc)-tdTomato fusion protein by binding to a TF-responsive promoter element. (C) IVIS images of TRE3GS-L2T mouse immortalized macrophages treated with LINQ1-tTA and the respective quantifications in (D) radiance and (E) fold change relative to the untreated control. (F) IVIS images of 5xUAS-L2T mouse immortalized macrophages treated with LINQ1-Gal4 and the respective quantifications in (G) radiance and (H) fold change relative to the untreated control. Area under the curve from the fold change graphs were quantified for (I) tTA and (J) Gal4. ****p<0.0001

## Results

### Quasiorganelles drive dynamic activation of a synthetic reporter circuit in mouse macrophages

We developed dynamic reporter systems in which synthetic organelles produce synthetic transcription factors that are secreted from the engineered organelles into the cytoplasm and are translocated to the nucleus of the host cell where they induce the transcription of a mammalian reporter gene, to create engineered interkingdom communication. Specifically, LINQ1 and *B. subtilis* were engineered to produce and secrete synthetic transcription factors originally derived from prokaryotic (tetracycline transactivator; tTA)^38^ and eukaryotic (Gal4) gene regulation pathways (Fig. 1A) ^38^. In parallel, a reporter immortalized bone marrow-derived mouse macrophage (iBMDM) cell line was engineered with a gene encoding a luciferase-tdTomato (L2T) fusion protein under the control of either a tTA- or Gal4-responsive promoter (Fig. 1B). This interkingdom reporter system was first validated by transient co-transfection into HEK293T cells of a plasmid expressing the transcription factor and the reporter construct, where we observed ∼200-fold induction after 48-h (Fig. S1A, B). Production and secretion of the synthetic transcription factors were then optimized in *B. subtilis* and LINQ1 by screening various signal peptides and N-terminal sequences that direct the protein for secretion (Fig. S2A, S3A,B). We observed that secretion was highest using the LLO and yoaW signal peptides in LINQ1 and *B. subtilis*, respectively.

The reporter system was then tested by infecting reporter macrophages with LINQ1 expressing tTA (Fig. 1C-E, I) or Gal4 (Fig. 1F-H, J). In both systems, there was a dose- and time-dependent increase in bioluminescence signals that peaked at 8-11-h post-infection (p.i.) at a multiplicity of infection (MOI) of 100 synthetic organelles per host cell. The peak fold induction was ∼100-fold using LINQ1-tTA and ∼80-fold using LINQ1-Gal4 (Fig. 1E, H; p<0.0001). While LINQ1-tTA had a higher dynamic range, LINQ1-Gal4 showed faster kinetics, with peak signal observed at 8-h p.i. (Fig. 1H) compared to 11-h p.i. for LINQ1-tTA (Fig. 1E). In contrast, reporter macrophages infected with *B. subtilis* LLO expressing tTA demonstrated only a ∼2-fold increase in bioluminescent signal that peaked at ∼16-h post-infection (Fig. S2B,C) at MOI-200. Given the versatility and safety of the auxotrophic *Listeria* and its greater dynamic range of gene induction we proceeded with LINQ1.

We further iterated on the system by testing the effect of growth conditions and the effect of dimerization of the synthetic transcription factor. LINQ1 expressing tTA was compared to LINQ1 expressing a single-chain dimer of tTA (sc-tTA), where two tTA monomers were connected by a (GGGGS)_4_ linker. Similar to previous results, we observed that LINQ1-tTA peaked at MOI-100 and reached ∼150-fold induction of bioluminescence (Fig. S4A,B). In contrast, LINQ1-sc-tTA only reached a peak induction of ∼50-fold, but the bioluminescence signal appeared to remain stable for longer than that of LINQ1-tTA (Fig. S4C,D). Growing LINQ1 cultures at 37°C instead of 30°C appeared to also have a significant effect on reporter gene induction. In these studies, peak induction was observed at MOI-10 (Fig. S4E-G; p<0.0001) and signal remained stable for longer than when LINQ1-tTA was grown at 30°C. In contrast to when LINQ1-tTA was grown at 30°C, we observed lower reporter gene induction at higher MOIs, indicating host toxicity likely caused by higher expression of genes such as listeriolysin O (LLO). Reporter curves correlated with intracellular bacterial counts determined at the end of the study (Fig. S4H, I).

The applicability of this reporter system across different cell types was demonstrated by testing in HEK293T cells. Unlike macrophages, these cells are non-phagocytic, and LINQ1 enters these cells by receptor-mediated endocytosis. HEK293T cells were transiently transfected with the reporter construct prior to infection with *Listeria* expressing tTA. Similar to studies in macrophages, a dose- and time-dependent increase in bioluminescence was observed with peak signal at MOI-10 (Fig. S5A,B). Since high plasmid copy number in HEK293T cells likely affects background levels and reporter gene induction, we engineered stable HEK293T cells expressing the L2T reporter under the control of a tTA- or Gal4-responsive promoter. LINQ1-tTA induced a 3-fold increase in luminescence signal at 24-h compared to controls (Fig. S5C,D). However, LINQ1-expressing tTA under the control of an intracellular-restricted promoter (pActA) did not produce a significantly higher luminescence signal (Fig. S5C,D). In contrast with LINQ1-tTA, LINQ1-Gal4 showed a ∼17-fold increase in bioluminescence signal at MOI-10 (Fig. S5E,F). In this case, there was significant induction of bioluminescence using pActA (Fig. S5E,F).

### Quasiorganelles reveal in vivo dynamics of transcriptional activation in a reporter mouse model

Since our in vitro results indicated that quasiorganelles activate transcriptional activation in mouse macrophages, we next tested the reporter system in a mouse model expressing luciferase under control of a tTA-responsive promoter (LC-1).^39,40^ The mouse model was validated by hydrodynamic transfection of a minicircle plasmid expressing tTA and by crossing the LC-1 mouse strain with a transgenic mouse engineered to express the reverse tetracycline transactivator (Fig. S1C,D).^41,42^ To understand spatial and temporal host-quasiorganelle dynamics in vivo, LC-1 and wild-type (WT) mice were injected intravenously with LINQ1-tTA and bioluminescence images were acquired over time (Fig. 2A,B). LC-1 mice had ∼25-fold higher induction of bioluminescence signal compared to WT mice. While in most cases signal lasted for 5-7 days (d) post-injection, some mice showed persistent signal for up to 24-d (Fig. 2A). Upon organ dissection on day 28 post-injection, the signal was largely localized to the lungs, spleen, and distal small intestine (Fig. S6A,B). This study was repeated, and organs were dissected after 24-h, where a similar distribution was observed with peak signal in the distal small intestine (Fig. S6C,D).

**Figure 2.**
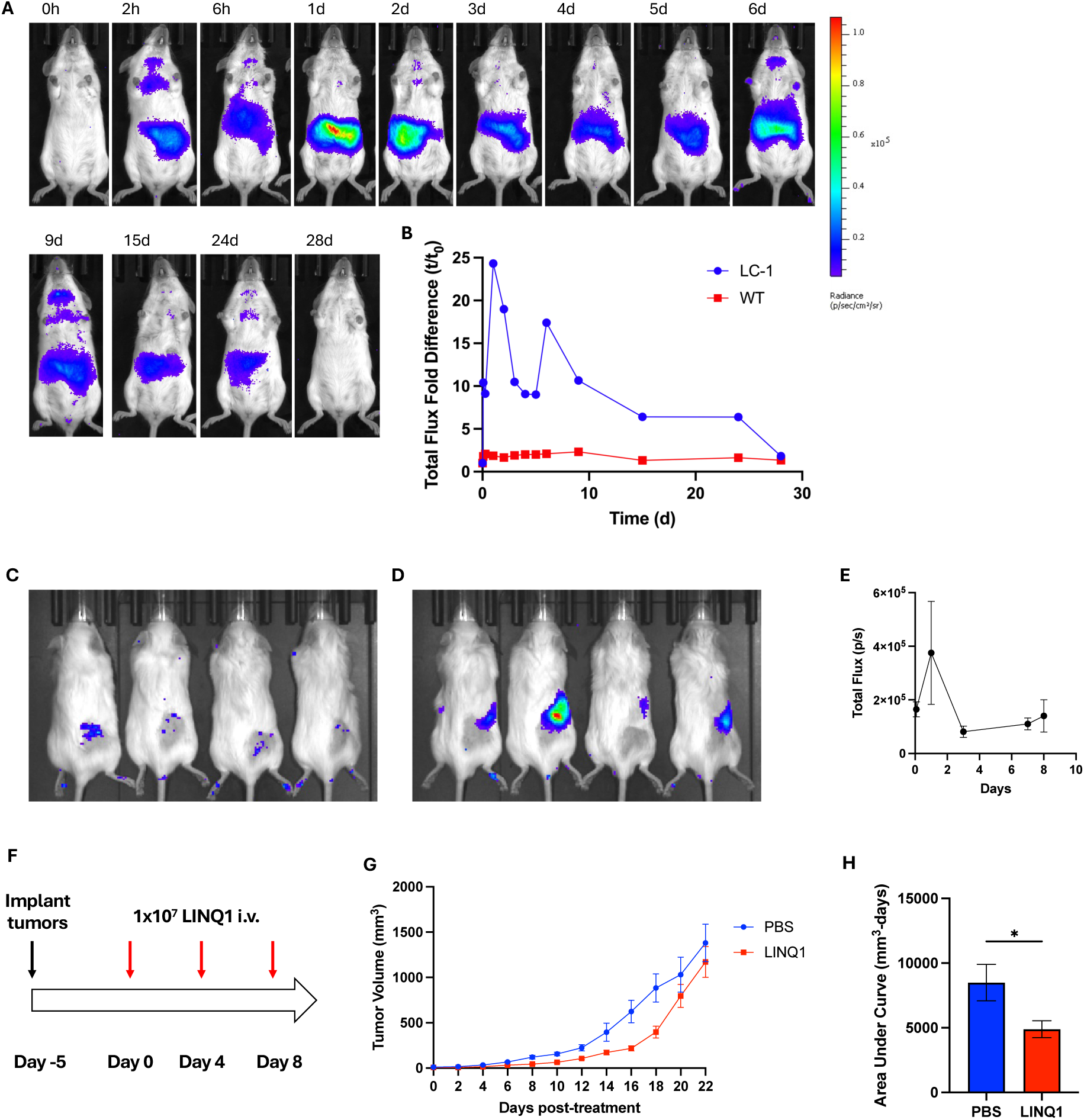
Dynamic activation of the reporter gene in LC-1 mice injected intravenously with LINQ1-tTA. (A) IVIS images of LC-1 mice injected with LINQ1-tTA over 28 d and (B) fold change of signal each day relative to the signal on day 0. (C) LC-1 mice implanted with subcutaneous E0771 tumors show activation at (C) 2 hours and (D) 24 hours post-treatment with LINQ1-tTA. (E) Kinetics and dynamics of E0771 tumor-bearing LC-1 mice injected with LINQ1-tTA, showing that tumors showed luminescence at 24 hours after the first injection but did not luminesce after the second injection. (F) Treatment regimen of orthotopic E0771 tumors in C57Bl/6 mice. (G) Quantification of tumor volume over 22 d and (H) quantified area under the curve. *p<0.05

We further tested this reporter model using the strain matched E0771 triple-negative breast cancer cells in LC-1 mice. Following intravenous injection of LINQ1, we observed ∼2-fold increase in bioluminescent signal, with significant variability between animals (Fig. 2C-E). Upon repeat injection, no significant activation was observed (Fig. 2E). The relevance of the reporter model was demonstrated in a therapeutic study, where C56Bl/6 mice were treated 3 times intravenously with LINQ1 or saline, and tumor volume was measured over time. Although we observed significant suppression of tumor growth, tumors eventually relapsed at 16-d post-treatment (Fig. 2F-H; p<0.05).

## Discussion

Controlling cellular fates and functions directly in living mammalian cells and mammals is a versatile approach with broad applications in tissue engineering, regenerative medicine, cancer therapy, and autoimmunity. Synthetic organelles, based on engineered intracellular microbes, are an ideal platform to achieve this goal for several reasons. Modeled after the origin of mitochondria, synthetic organelles can be engineered to produce mammalian proteins directly inside mammalian cells, and these can be targeted to the nucleus or directed to control cytoplasmic signaling networks. Bacteria-expressed transcription factors can be used to reprogram mammalian functions, targeting key nodes in cellular gene circuits, and similarly to control cellular function targeting cytoplasmic regulatory pathways.

Synthetic organelles benefit from the rules of microbial genetic design using polycistronic operons, where a single genetic promoter element can drive the expression of multiple downstream genes, including mammalian regulatory proteins when encoded in EES as operons. The respective proteins can be designed to affect given regulatory networks using multiple control points or multiple pathways simultaneously, either in the cytoplasm or within the nucleus. In addition, promoters in EES can be controlled orthogonally to mammalian genes using various exogenous stimuli including heat,^43^ light,^44^ ultrasound,^45^ small molecules,^46^ or using autonomous promoters that respond to population density or specific stimuli like pH, hypoxia, and lactate.^47–49^ Such control strategies can be used to direct mammalian cell function from a distance, e.g. from outside the bodey. Lastly, in comparison with alternative methods to achieve cellular reprogramming in vivo, synthetic organelles do not modify the genome of the host cell, and thus pose a lower risk for oncogenesis due to gene insertion, and can be eliminated readily using readily available kill-switches or antibiotics.

Cancer is a heterogenous family of diseases driven by genetic and environmental factors, and therefore, single-agent therapies targeting individual molecular drivers of malignancy have proven inadequate. Cancer cells develop escape mechanisms by mutation, clonal selection, transcriptional and metabolic reprogramming, and by modulating the TME. Overall, these changes lead to therapeutic resistance and subsequent disease recurrence. The TME, composed of interstitial cells, such as fibroblasts and endothelial cells, and immune cells, such as T cells and macrophages, contributes significantly to therapy resistance, but can share a number of common features. Previously, we demonstrated that synthetic organelles based on *B. subtilis* could alter mammalian cell phenotypes for enhanced anti-tumor effects in triple-negative breast cancer.^1,2^ In these studies, the synthetic organelles produced transcription factors to modulate the phenotype of macrophages towards pro-inflammatory or anti-inflammatory fates.^1,2^ However, a key limitation was that *B. subtilis*-based synthetic organelles were eliminated, likely via autophagy, by 12-h post-infection in primary mouse macrophages.^1,2^ Thus, it was necessary to develop these quantitative systems to measure interkingdom communication between synthetic organelles and host cells, providing a foundation for understanding and ultimately improving their intracellular persistence and therapeutic functionality.

Here, we describe a reporter system that uses two synthetic transcription factors, tTA and Gal4, to activate expression of luciferase within host cell sin culture and in animals as a way to model interkingdom communication for cellular control. This system revealed the magnitude, kinetics, and spatial distribution of LINQ1-based synthetic organelles and demonstrated that LINQ1 can control the expression of genes within a mammalian host cell. In addition, this reporter circuit enables optimization of various parameters that could influence communication between synthetic organelles and host cells. Between the two chassis organisms used for the development of synthetic organelles, the *Listeria*-based LINQ1 chassis activated host cells ∼100-fold compared to ∼2-fold with *B. subtilis*. This reflects cumulative differences between these chassis in terms of infectivity, avoidance of autophagy, protein secretion, and proteolytic cleavage of engineered proteins. *B. subtilis* produces many extracellular proteases, and these proteases are often knocked out in strains used for protein production.^50^ However, we observed that when these *B. subtilis* strains are engineered to express multiple mammalian transcription factors, the resulting strains are genetically unstable and mammalian proteins still undergo cleavage in these strains, indicating the presence of additional mechanisms of proteolytic cleavage. In contrast, we observed higher protein secretion and lower rates of proteolytic cleavage in LINQ1, which likely enables higher activation of host cells.

The dynamic range of reporter gene activation in vivo is likely limited by its location in the genome, which is not fully characterized in the LC-1 mouse model.^39^ Immune cells like macrophages and lymphocytes typically have lower expression of reporter genes compared to non-immune cell types, and gene silencing occurs frequently in these cells.^51^ Future studies could use the Gal4 reporter mouse model to determine the differences in spatial distribution and dynamic activation using synthetic organelles in animals.^52^

Lm has been developed extensively as an anti-cancer immunotherapy,^27–35^ largely using the live-attenuated double-deleted (LADD; ΔactA ΔInIB) mutant or the D-alanine auxotroph (Δdal Δdat) mutant. In tumors, Lm primarily infects phagocytic immune cells such as TAMs, tumor-associated dendritic cells, and myeloid-derived suppressor cells (MDSCs),^53^ but it can also directly infect tumor cells depending on the tumor type.^54^ Intravenous administration of Lm vaccine strains in tumor-bearing mice has been shown to increase intratumoral CD8^+^ T cells and antigen-presenting dendritic cells, reprogram intratumoral MDSCs and TAMs to a pro-inflammatory phenotype, and reduce intratumoral regulatory T cell populations.^53,55–57^ However, clinical translation of Lm-based therapies has been restricted by safety concerns due to extracellular survival of commonly used strains.^58,59^

To address these concerns and facilitate clinical translation, the chassis used in this study was engineered with FAD and FMN auxotrophies and deletions in *actA* and *inIB*, largely restricting its distribution to phagocytic immune cells and eliminating extracellular reservoirs, unlike wild-type *Listeria monocytogenes,* which has significant populations in the gallbladder, spleen, and liver of mice.^22,23,58,59^ In transgenic mice injected with LINQ1-tTA, we observed localization of bioluminescent signal in the submandibular lymph nodes, lungs, distal small intestine, and spleen. These results parallel the biodistribution of bioluminescent *Listeria*, although higher uptake in the liver is observed in these studies due to the expression of *inIB*. We also observed activation of luminescence in the tumor of reporter mice injected with LINQ1-tTA, although this signal was transient and refractory to repeated injections. In conjunction with the reporter data, these studies indicate that aggressive breast tumors evolve to resist quasiorganelle therapy, likely by restricting infiltration of myeloid immune cells into the TME.

Engineered intracellular microbes are emerging as a tool to redirect host cell biology in eukaryotic organisms. Modeled after the origin of mitochondria and chloroplasts,^60^ these systems typically use free-living organisms to transfer specific functions into fungal or mammalian host cells. The first challenge in developing synthetic organelles is uptake: microbes need to be engineered to express uptake proteins^1,2,61^ or are coated with polymers^62–68^ to enable uptake into host cells. In some studies, this challenge is bypassed by directly injecting the microbes into the host cell.^69,70^ The second major challenge is intracellular persistence and avoiding clearance. In the process of entering host cells, microbes can trigger endosomal and cytosolic sensing pathways like toll-like receptors, nod-like receptors, and cGAS-STING, which activate anti-microbial responses in the host cell. Using host-adapted intracellular organisms like *L. monocytogenes* or BCG is a possible solution, as these naturally avoid detection and clearance in host cells.^36^

The third major challenge is driving long-term endosymbiosis. In some studies, long-term persistence is forced by using auxotrophic host cells where the synthetic organelle rescues auxotrophy by producing a protein or metabolite. However, free-living bacteria have significantly faster growth rates compared to eukaryotic cells, which leads to host cell lysis due to overgrowth of bacteria. Using chassis organisms with minimized genomes like JCVI-syn3A^71,72^ or host-adapted organisms with slower growth rates like BCG could enable long-term symbioses, but these organisms are less genetically tractable due to their naturally slower growth rates. Engineering inheritance of synthetic organelles using surface display of mitochondrial proteins such as Miro may enable long-term persistence by segregating organelles into daughter host cells during cell division. For each of these challenges, a major limitation in the field is the lack of a reporter system to refine each variable independently to optimize long-term endosymbiosis. The reporter system developed here, where the synthetic organelle produces a synthetic transcription factor to activate luminescence in host cells, fills this gap to enable rapid optimization of variables and works across different chassis and host cell types.

Overall, the work described here provides a way to characterize the communication mediated by synthetic quasiorganelles in both in vitro and in vivo models. We characterize the transcriptional activation in response in host cells to two chassis organisms, *Listeria monocytogenes* and *Bacillus subtilis*, which demonstrated significant differences, highlighting the importance of chassis selection. Although our findings demonstrate the potential of quasiorganelles as cancer therapeutics, limitations remain in achieving sustained intracellular persistence and therapeutic efficacy following repeated administration. By enabling quantitative characterization of quasiorganelle-mediated transcriptional activity, the reporter system provides a framework for guiding the engineering of more effective quasiorganelle therapies.

## Materials and Methods

### Bacterial strains and growth conditions

*L. monocytogenes* and *E. coli* strains are listed in Supplementary Table 1. *E. coli* and *L. monocytogenes* were grown in Luria-Bertani (LB) and brain heart infusion (BHI) broth, respectively. For selection in *E. coli*, antibiotics were used at the following concentrations: chloramphenicol (25 µg/ml), carbenicillin (100 µg/ml), kanamycin (50 µg/ml). For selection in *L. monocytogenes*, antibiotics were used at the following concentrations: chloramphenicol (10 µg/ml), erythromycin (1 µg/ml), kanamycin (50 µg/ml), nalidixic acid (50 µg/ml), streptomycin (200 µg/ml). *E. coli* and *L. monocytogenes* strains were kind gifts from Drs. Daniel Portnoy, John-Demian Sauer, and Darren Higgins. The Δ*ribC* Δ*ribF* strain was grown in BHI broth or agar containing FAD or FMN as described previously.^22,23^

### Mammalian cell culture

Mammalian cells were grown in Dulbecco’s modified Eagle medium (DMEM; Gibco) with 10% (vol/vol) fetal bovine serum (FBS; Gibco) at 37°C and 5% CO_2_ unless otherwise specified. Cells were subcultured by routine culture protocols and detached using TrypLE Express (Gibco). Cells were tested periodically for mycoplasma and confirmed to be mycoplasma negative.

### Plasmid cloning

The backbone DNA was amplified using 2x Phusion polymerase master mix (ThermoFisher) and run on a thermocycler per manufacturer’s recommendations. PCR fragments were run on a 0.6% agarose gel (Fisher Scientific) at 100V for 30 minutes, and purified using a gel extraction kit (New England Biolabs), after which their concentration was measured on a NanoDrop (Fisher Scientific). For bacterial constructs, synthetic transcription factor genes (tTA, Gal4-VP16, sc-tTA) were synthesized as gene fragments (Twist Bioscience). The gene fragment and backbone were mixed at a molar ratio of 3:1 with 2x Hi-Fi master mix (New England Biolabs) using 50 ng of backbone and incubated at 50°C for 60 min. The final product was transformed into NEB5α or NEB Turbo (New England Biolabs) as recommended by the manufacturer. Colonies were subsequently inoculated in LB at 30°C or 37°C before isolation via a plasmid miniprep kit (Qiagen). The final construct was validated both by running a restriction digest and DNA sequencing (Plasmidsaurus). Lentivirus backbone was amplified from pLenti CMV Puro LUC (w168-1) (Addgene plasmid #17477) using 2x Phusion polymerase master mix (ThermoFisher) and manufacturer’s recommendations. The tetracycline-responsive element (TRE3) promoter was amplified from pTRE3G (Takara Bio) and the luciferase-tdTomato fusion gene was amplified from pcDNA3.1(+)/Luc2=tdT (Addgene plasmid #32904) using 2x Q5 polymerase master mix (New England Biolabs) and manufacturer’s recommendations.

### Minicircle Plasmid Generation

Minicircle plasmid backbone (pMC.BESPX-rtTA) was transformed into competent *E. coli* ZYCY10P3S2T (System Biosciences; MN900A-1). On day 1, a 2 mL starter culture was inoculated in LB containing kanamycin and grown at 37°C 250 rpm overnight. The following day, 50 µL of the starter culture was inoculated into 200 mL of Terrific Broth (Invitrogen; 22711022) with kanamycin and grown at 37°C 250 rpm overnight. The next day, optical density of the culture was measured and confirmed to be in the range of 4-6 prior to minicircle production. 200 mL of minicircle induction mix was prepared in a separate flask using 192 mL LB, 8 mL 1N sodium hydroxide, and 200 µL of fresh 20% L-arabinose. The 200 mL culture was mixed with 200 mL induction mix and grown at 32°C 250 rpm for 5 hours. Bacterial pellets were isolated by centrifugation and kept at -20°C until purification. Plasmids were purified using the ZymoPURE II plasmid maxiprep kit (Zymo Research; D4202) following the manufacturer’s protocol for endotoxin-free purification.

### SM10 λpir *E. coli* transformation

*E. coli* SM10 λpir glycerol stock was streaked on an LB agar plate containing 50 µg/mL kanamycin and grown at 37°C overnight. The next day, bacteria were inoculated into 20 mL of LB and grown at 37°C in a shaking incubator set to 225 rpm for 2 hours. Bacteria were centrifuged at 4,500 g for 10 min, and the supernatant was discarded. The pellet was resuspended in 1 mL of ice-cold 100 mM CaCl_2_ and placed on ice. 100 µL of bacteria was aliquoted into a 1.5 mL microcentrifuge tube and 1 µg of plasmid DNA was added. After 30 min on ice, the tube was placed at 42°C for 90 sec. After another 5 min on ice, 900 µL of SOC broth (NEB) was added and the bacteria were incubated at 37°C in a shaking incubator set to 225 rpm for 1 hour. The tube was centrifuged at 12,000xg for 3 min, and 900 µL of supernatant was discarded. The pellet was resuspended in the remaining 100 µL and spread on an LB agar plate containing 25 µg/mL chloramphenicol. Colonies were verified by PCR using 2x Phire polymerase master mix (ThermoFisher) and manufacturer’s recommendations.

### Conjugation of plasmid into *Listeria monocytogenes*

Donor *E. coli* SM10 λpir containing the plasmid of interest was streaked on a BHI agar plate and incubated at 30°C. Separately, *Lm* was streaked onto a BHI agar plate supplemented with 10 µL each of FAD and FMN, and incubated overnight at 37°C. The following day, SM10 was streaked on a BHI agar plate without antibiotics with 10 µL each of FAD/FMN in a circular pattern and *Lm* was streaked directly over the *E. coli* in the same circular pattern. The plate was incubated at 30°C overnight. The next day, a patch of growth was collected using a loop and resuspended in 500 µL of sterile PBS. 100 µL of undiluted solution and 100 µL of 1:10 and 1:100 dilutions were plated on BHI plates with appropriate antibiotics and FAD/FMN. The plates were incubated at 30°C for 1-d and then 37°C for 1-d. Colonies were visible after 2-d. 4-6 colonies were re-streaked on a new plate, genotyped by PCR, and then inoculated into BHI broth with antibiotics and FAD/FMN. Cultures were grown in a shaking incubator at 225 rpm and 37°C for 18-h. Glycerol stocks were made by adding 500 µL of 50% sterile glycerol:BHI and 500 µL liquid culture, and stored at -80°C.

### Phage Transduction

Phage transduction was performed based on previously published protocols.^73^ To prepare phage lysate for transduction, donor *Lm* was grown at 30°C in LB broth in a stationary incubator. The next day, sterile top agar (LB + 0.7% agar) was microwaved, aliquoted in 14 mL conical tubes in 4 mL per tube, and cooled to 55°C in a water bath. 1 mL of donor culture was spun down and resuspended in 1 mL LMC broth (LB + 10 mM MgSO_4_ + 10 mM CaCl_2_). MgSO_4_ and CaCl_2_ were added to the tube of top agar to a final concentration of 10 mM, along with 400 µL of donor culture in LMC, and 10 µL of U153 phage stock. The tube was mixed quickly by swirling, poured on a BHI agar plate, and incubated at 30°C overnight. The next day, 5 mL of sterile TM buffer (10 mM Tris-HCl, 10 mM MgSO_4_ in MilliQ water) was added to the plate and incubated at 4°C overnight. The next day, the liquid was transferred to a new 15 mL tube and filtered through a 0.22 µm filter. This phage lysate was stored at 4°C indefinitely, and 100 µL was plated on a BHI agar plate to confirm sterilization. For phage transduction, recipient *Lm* was grown at 30°C in LB broth in a shaking incubator. The next day, the concentration of *Lm* was adjusted to 5x10^8^ CFU/100 µL in LMC broth, and 10-20 µL of phage lysate was added. The volume was adjusted to 200 µL and incubated at room temperature for 30 minutes. The entire mixture was plated on a BHI agar plate with appropriate antibiotics and placed at 37°C. Colonies were visible after 2-3 days. Transductants were restreaked on fresh plates with appropriate antibiotics and genotype was confirmed by PCR.

### Western blot sample preparation for secretion screening

To assess protein secretion by *Lm*, bacterial cultures were grown overnight in BHI broth at 37°C with shaking at 200 rpm. The following day, 1 mL of each culture was collected and centrifuged at 16,000 xg for 3 min at room temperature. A 75 µL aliquot of the resulting supernatant was transferred to a 1.5 mL microcentrifuge tube and combined with 4× Laemmli sample buffer (Bio-Rad) supplemented with 1.42 M 2-mercaptoethanol (Bio-Rad). Samples were briefly vortexed and heated at 95°C for 15 min in a thermocycler (Thermo Fisher Scientific), followed by cooling to room temperature. For SDS-PAGE, 10 µL of each prepared sample was loaded onto a 4–20% polyacrylamide gel (Bio-Rad) alongside 5 µL of protein ladder. Gels were electrophoresed at 100 V for 80 min. Proteins were subsequently transferred to a PVDF membrane (Bio-Rad) using a semi-dry transfer system at 25 V for 30 min. Following transfer, membranes were blocked for 1-h at room temperature with gentle rocking in blocking buffer (Bio-Rad). Membranes were then incubated overnight at 4°C with rocking in mouse monoclonal anti-FLAG primary antibody (BioLegend, 637315) diluted in 1× Tris-buffered saline containing 0.1% Tween-20 (TBS-T) and 1% bovine serum albumin (BSA). The following day, membranes were washed three times in TBS-T for 10 min per wash and incubated for 1-h at room temperature with gentle rocking in HRP-conjugated anti-mouse secondary antibody (Cell Signaling Technology, 7076S) diluted in TBS-T containing 1% BSA. After three additional 10-min washes in TBS-T, membranes were incubated with SuperSignal West Femto chemiluminescent substrate (Thermo Fisher Scientific) for 5 min. Chemiluminescent signal was then detected using a ChemiDoc MP imaging system (Bio-Rad).

### Mouse strains

The LC-1 C57Bl/6 mouse strain was a kind gift from Dr. Marcus Mall.^40^ This strain expresses firefly luciferase and cre recombinase under control of a bidirectional tetracycline responsive element (TRE) promoter. LC-1 mice were crossed to C57Bl/6 albino mice (Jackson Laboratories; strain: 000058) for several generations to obtain mice with a white coat for in vivo bioluminescence imaging (BLI). Mice were genotyped as indicated and bred to homozygosity. LC-1 mice were crossed to B6.Cg-*Gt(ROSA)26Sor^tm1(rtTA*M2)Jae^*/J (Jackson Laboratories; strain: 006965) for validation studies. These mice were placed on a doxycycline diet composed of 625 mg/kg doxycycline hyclate (Inotiv; TD.01306) for 7-d, and then injected with 100 µL of D-luciferin by intraperitoneal injection (i.p.) and imaged on an in vivo imaging system (IVIS).

### Mouse genotyping

Mouse tail samples were obtained at weaning and stored at -20°C. Genomic DNA was extracted using the Monarch Spin gDNA extraction kit (New England Biosciences; T3010S) according to manufacturer’s protocol for tissue samples. Extracted genomic DNA were amplified using polymerase chain reaction (PCR) for the LC-1 transgene using Phire Hot Start II PCR master mix green (Thermo Scientific; F126S) according to manufacturer’s protocol. PCR samples were run on a 1% agarose gel in 1x tris-acetate-ethylenediaminetetraacetic acid (TAE) buffer with Sybr Safe DNA gel stain (Thermo Scientific; S33102). Gels were imaged on a BioRad Chemidoc Touch Imaging System.

### Hydrodynamic transfection

10 µg of plasmid DNA was diluted in ∼2 mL PBS and injected by tail vein in 5 seconds or less. Mice were imaged at indicated time points using an IVIS following administration of 100 µL of D-luciferin by intraperitoneal injection (i.p.).

### Lentivirus preparation and transduction

Cells were seeded in 12-well plates at 2 × 10⁵ cells per well one day prior to lentiviral transduction. On the day of transduction, lentiviral stocks were thawed on ice. Culture medium was replaced with 400 µL of fresh DMEM containing µg/mL polybrene. Lentiviral stocks were prepared in DMEM containing polybrene to a final volume of 100 µL per well. To evaluate transduction conditions, varying volumes of lentiviral stock (10, 15, 25, or 50 µL) were combined with 90, 85, 75, or 50 µL of DMEM, respectively. The prepared lentiviral solutions were subsequently added to the corresponding wells, resulting in a total culture volume of 500 µL per well. Cells were incubated with lentivirus for 24-h under standard culture conditions. At 24-h post-transduction, lentivirus-containing medium was removed and replaced with fresh culture medium. Cells were monitored for confluency and, when appropriate, passaged into 6-well plates. Puromycin selection was initiated approximately 24-h after passaging. Prior to selection, a puromycin kill curve was performed using concentrations ranging from 1-12 µg/mL to determine the appropriate selection concentration for the cell line. Following initiation of puromycin selection, cells were maintained in puromycin-containing medium and passaged as necessary.

### In vitro infections with *Listeria monocytogenes* quasiorganelles

*Lm* constructs were cultured overnight in brain heart infusion (BHI) broth supplemented with 0.1% FAD and FMN at 37°C and shaking at 200 rpm. Overnight cultures were subsequently subcultured into fresh BHI broth containing 0.1% FAD and FMN at a 1:12 culture-to-broth volume ratio and grown under the same conditions until reaching logarithmic phase (OD₆₀₀ = 0.6–1.0). Bacterial concentrations were estimated from OD_600_ measurements using the approximation that an OD_600_ of 1.0 corresponds to 1 × 10^9^ *Listeria*/mL. The appropriate volume of bacterial culture required to achieve each multiplicity of infection (MOI) was calculated and aliquoted separately. Each bacterial aliquot was combined with DMEM to prepare an infection inoculum with a final volume of approximately 200 µL per well. Target cells were seeded in 24-well plates at 5 × 10^4^ cells per well one day prior to infection. Immediately before infection, the existing culture medium was aspirated and replaced with 200 µL of the prepared DMEM-containing bacterial inoculum corresponding to the desired MOI. Cells were co-incubated with *Lm* for 30 min, after which the inoculum was removed, and the wells were washed three times with PBS. Fresh culture medium was then added. At 30 min following medium replacement, gentamicin was added to a final concentration of 10 µg/mL to eliminate remaining extracellular bacteria. For longitudinal bioluminescence imaging, plates were imaged hourly using an IVIS imaging system. D-luciferin was added at 10 µL/mL prior to imaging, as appropriate.

### LC-1 reporter studies with subcutaneous E0771 mouse breast cancer implantation

6-8-week-old female (n=4) C57Bl/6 albino LC-1 mice^39,40^ were used for in vivo reporter studies. Mice were placed under anesthesia using isoflurane and then fur was removed using clippers. 2 x 10^5^ E0771 mouse breast cancer cells were implanted subcutaneously using a 27G needle attached to a 1 mL syringe. When tumors reached ∼50 mm^3^, mice were injected intravenously with 100 µL of 1 x 10^7^ CFU of LINQ1 expressing tTA by tail vein injection. Injections were repeated 7-d after the first injection. 100 µL of D-luciferin was administered by i.p. injection and mice were imaged on an IVIS at the indicated time points.

### Therapeutic studies using LINQ1 in E0771 orthotopic tumor model

6-8-week-old female (n=6-9) C57Bl/6 albino mice (Jackson Laboratories; strain: 000058) were used for tumor studies. Mice were placed under anesthesia using isoflurane and abdominal fur was removed using clippers and application of Nair for 30 seconds. 2 x 10^5^ E0771 mouse breast cancer cells were implanted subcutaneously using a 27G needle attached to a 1 mL syringe. Tumors reached 5 mm^3^ after 5-d. Mice were anesthetized with isoflurane prior to administration of quasiorganelles. Mice were injected with 1 x 10^7^ CFU of LINQ1 in 100 µL or 100 µL of saline by retro-orbital injection. Following injection, mice were monitored until recovery from anesthesia before being returned to their home cages. Injections were repeated every 4-d for a total of 3 injections. Tumor volume was measured at the indicated time points using calipers and volume was calculated using the formula: 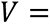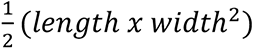

### Colony forming unit quantification

Macrophages were infected as indicated and processed after the final imaging time point to collect intracellular bacteria for quantification.^74^ Media was aspirated and replaced with lysis buffer (milliQ water, 10 mM Tris, 0.1% Triton X-100) and rocked gently for 5 min. Lysates were diluted in PBS and plated onto BHI agar plates with FAD and FMN. Colonies were visualized after 2-d of incubation at 37°C.

## Statistical analysis

Statistical analyses were performed using the GraphPad Prism software (version 11.0.1). Statistical tests are indicated for each experiment. Data were expressed as mean ± standard deviation, using 3 or more biological replicates for each group. Threshold for significance was set as p < 0.05 and p values were adjusted for multiple comparisons depending on the statistical test used.

## Data Availability

Raw data and bacterial strains will be made available upon reasonable request to the corresponding author.

## Acknowledgments

The authors thank Dr. Daniel Portnoy, Dr. John-Demian Sauer, and Dr. Darren Higgins for the *Listeria monocytogenes* strains and plasmids used in this study. The authors thank Dr. Michael Bachmann for reporter gene plasmids used in this study. The authors thank Dr. Marcus Mall and Dr. Julia Duerr for the LC-1 mouse strain. The authors thank Dr. Andrew Olive for the iBMDM cell line.

---

**Figure S1.**
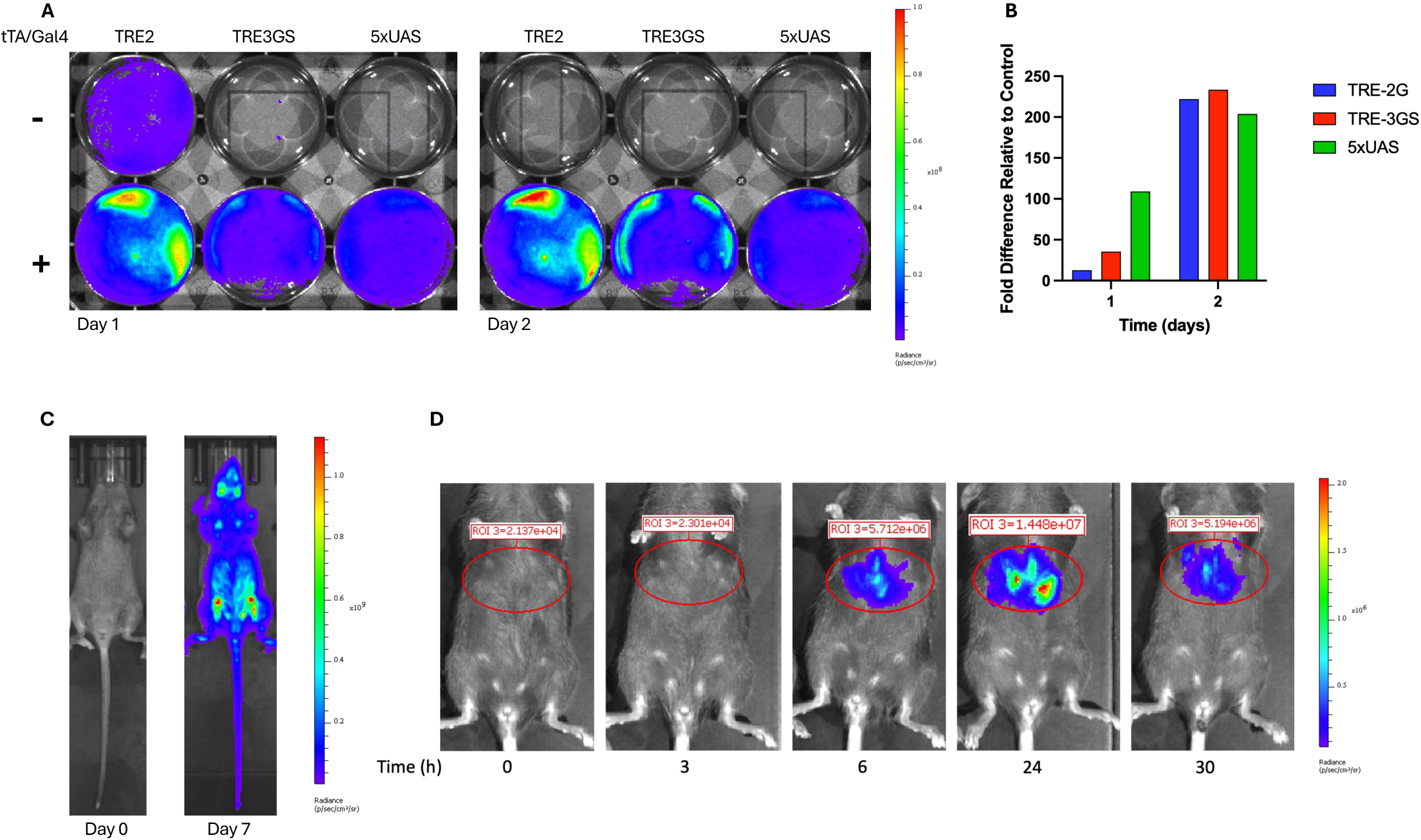
Validation of reporter systems in culture and mouse models. (A) HEK293T cells transiently transfected with the reporter construct and the corresponding synthetic transcription factor. (B) Activation of the reporter gene relative to the control that did not receive the synthetic transcription factor construct. (C) LC-1 mice crossed to R26-M2rtTA and fed doxycycline for 7 d prior to imaging on the IVIS optical system. (D) IVIS images of LC-1 mice transfected in vivo by hydrodynamic injection of a minicircle plasmid encoding tTA.

**Figure S2.**
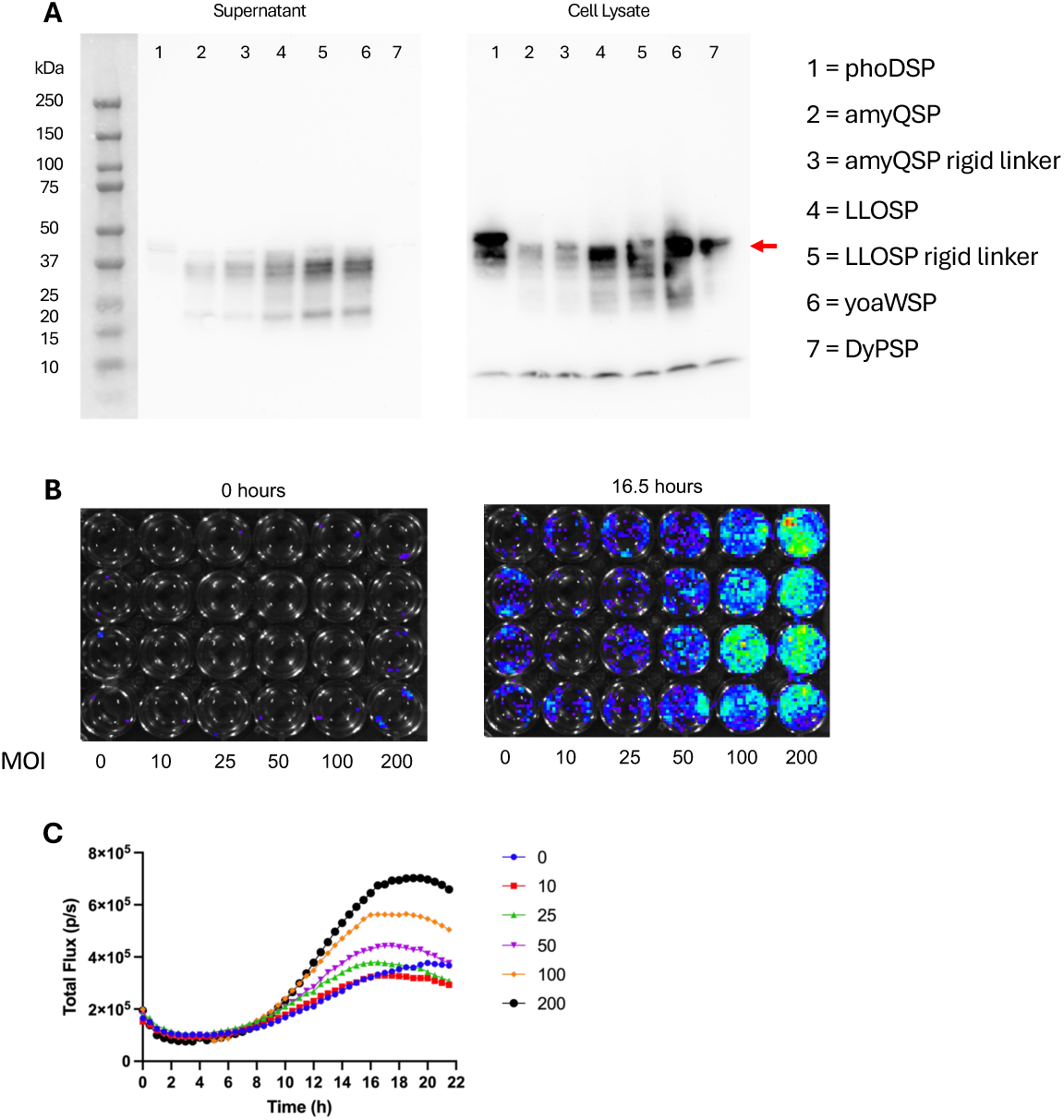
Validation of the reporter system in *B. subtilis*-LLO. (A) Secretion study of tTA with various signal peptides derived from *B. subtilis* or *L. monocytogenes* comparing supernatant and cell lysate samples. The red arrow indicates the expected size of the protein. (B) IVIS images of TRE3GS-L2T mouse immortalized macrophages treated with *B. subtilis*-tTA. (C) Kinetics and dynamics of reporter gene in mouse immortalized macrophages treated with B. subtilis-tTA, with peak signal at ∼16.5 hours post-treatment.

**Figure S3.**
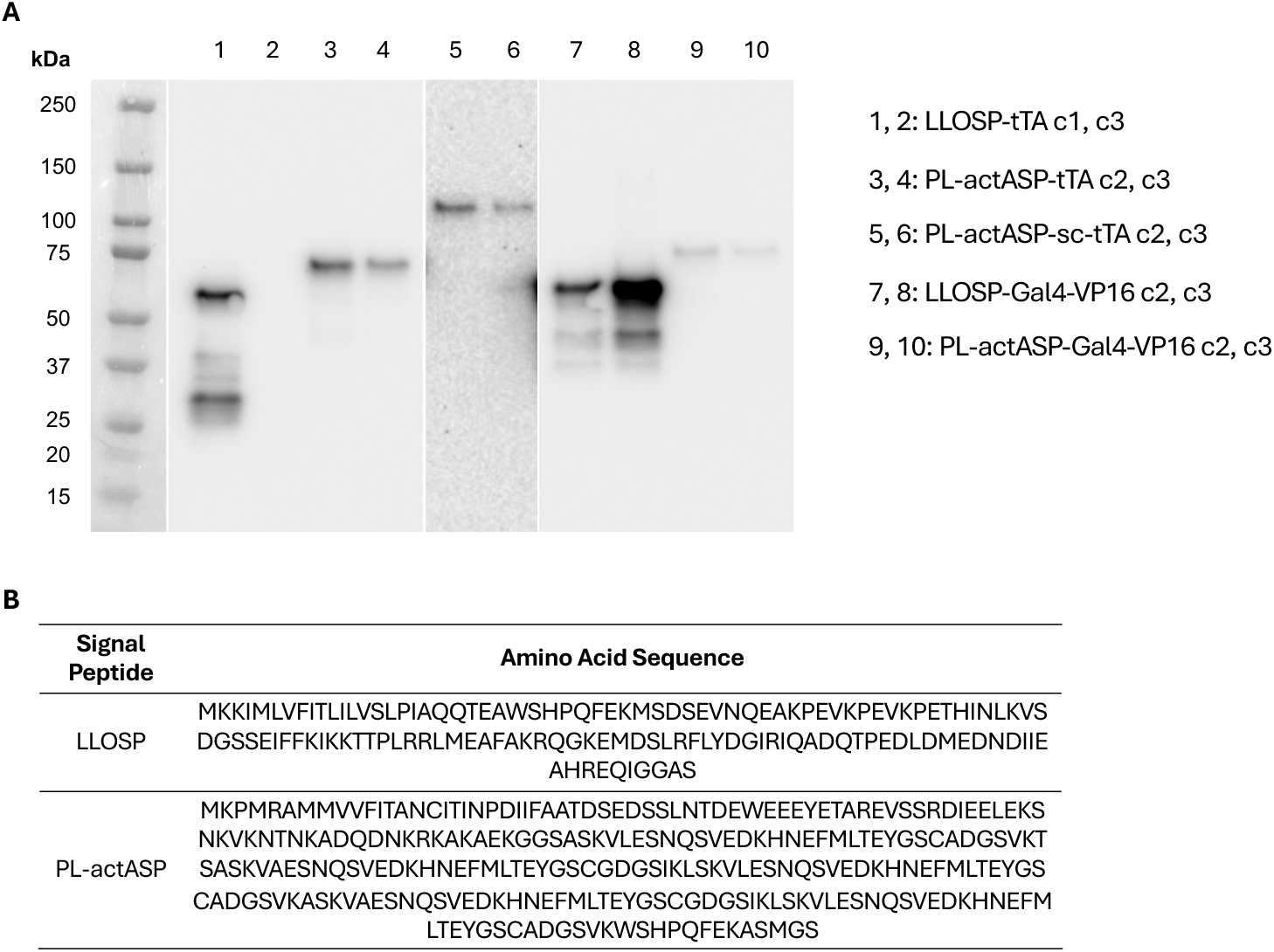
Secretion study of tTA and Gal4 in *Listeria monocytogenes*. (A) Western blot of supernatants with various signal peptides for tTA, single-chain dimeric tTA, and Gal4 with (B) the respective sequences of the signal peptides.

**Figure S4.**
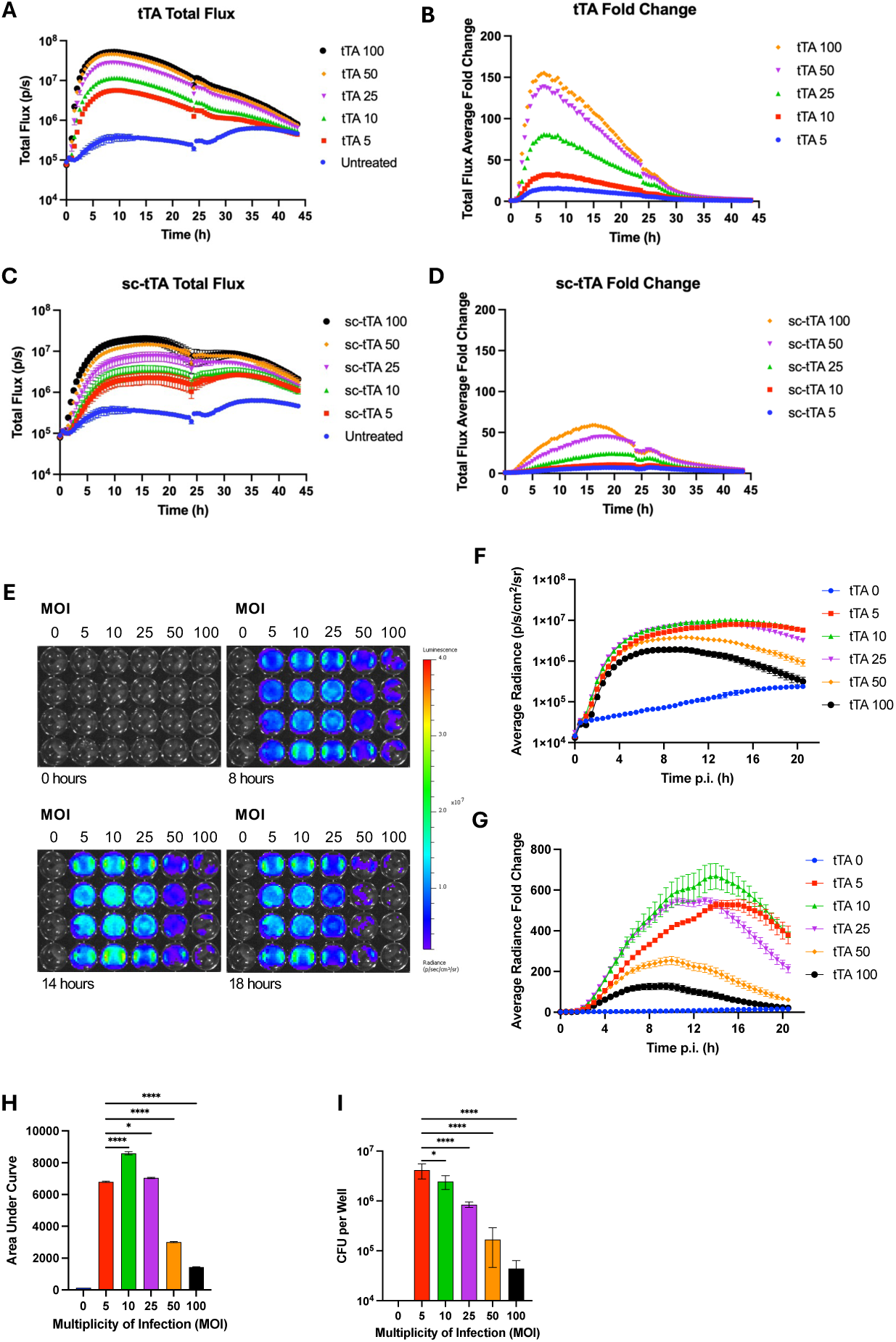
Refining the host-quasiorganelle relationship by modulating transcription factor stability and quasiorganelle growth conditions. Comparing activation of the reporter gene in TRE3GS-L2T immortalized mouse macrophages treated with (A, B) LINQ1-tTA or (C, D) LINQ1-sc-tTA shows differential kinetics and dynamics of reporter gene activation. The effect of growing LINQ1 at 37°C at multiple MOIs was assessed in (E) IVIS images, (F) radiance counts, and (G) fold change relative to signal at time zero (t0). The (H) quantified fold change graph parallels the (I) quantified colony-forming units of bacteria. *p<0.05, ****p<0.0001

**Figure S5.**
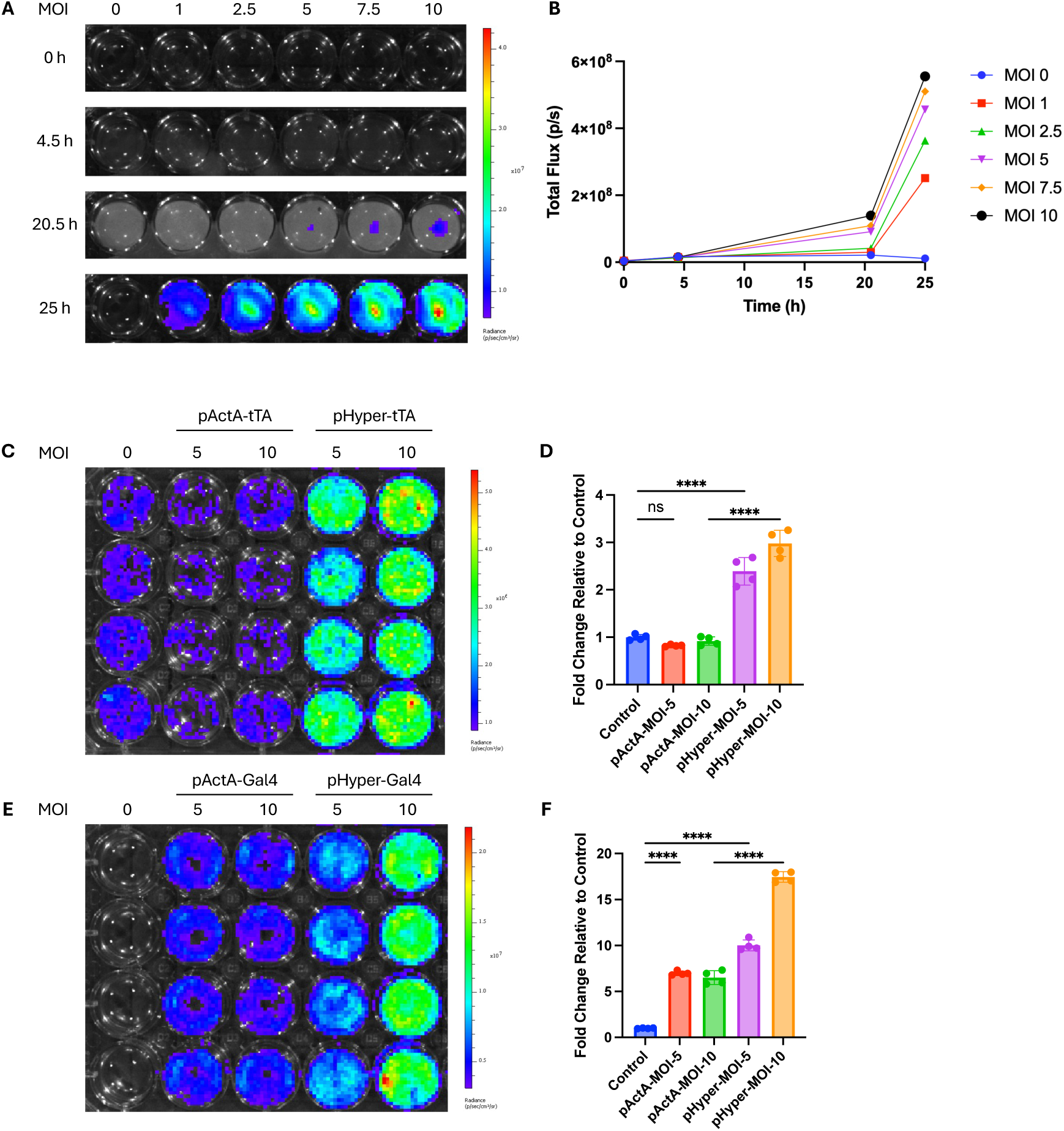
Activation of the reporter gene circuit in non-phagocytic cells. HEK293T cells were transiently transfected with the reporter construct and then infected 24 h later with LINQ1-tTA. Images were taken on the IVIS optical system at various time points (A) and were quantified (B). Reporter gene induction was assessed in HEK293T cells stably expressing (C, D) TRE3GS or (E, F) 5xUAS construct, using either a constitutive promoter (pHyper) or the intracellular-activated promoter (pActA). ****p<0.0001

**Figure S6.**
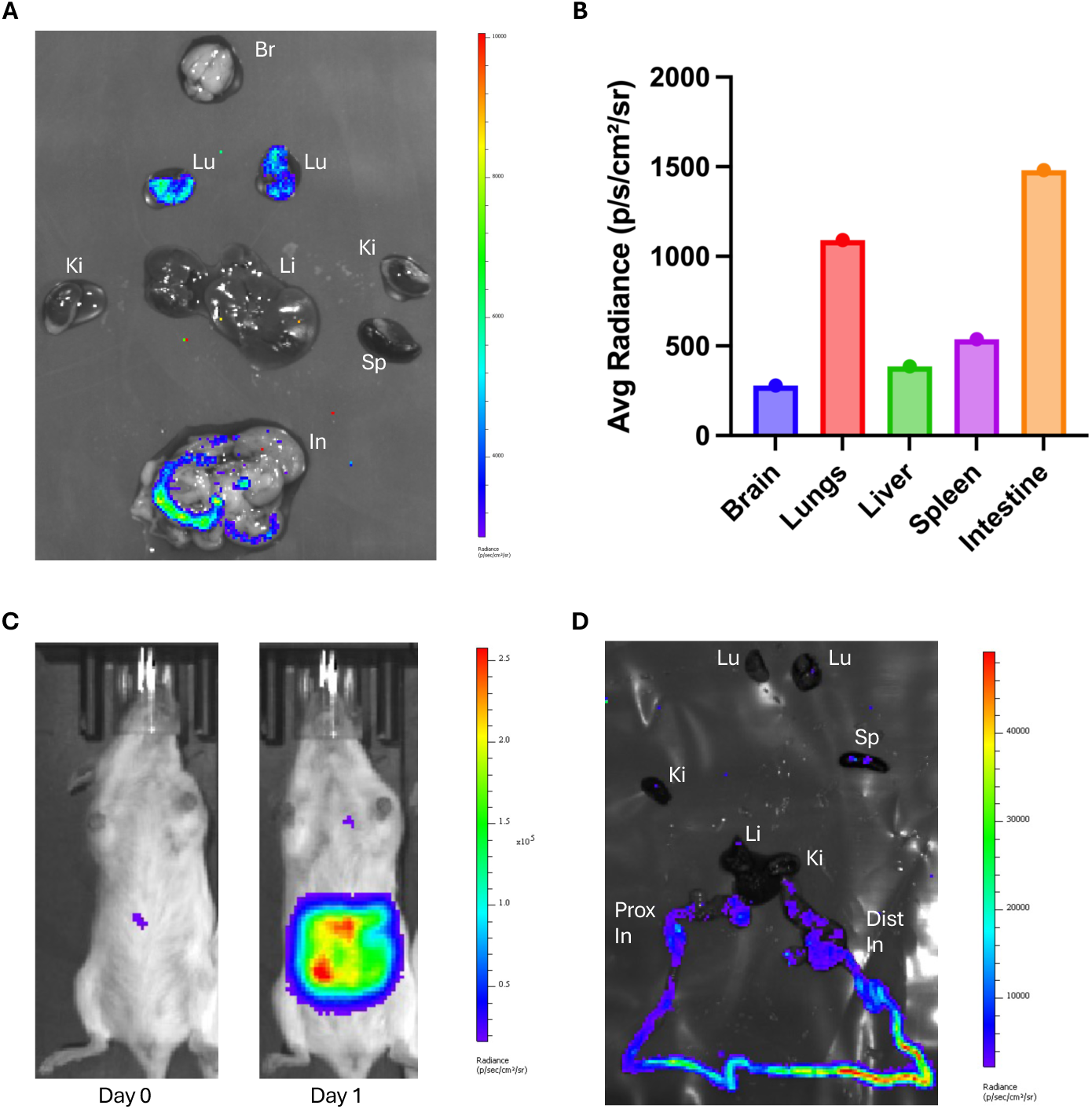
Distribution of quasiorganelles in vivo. LC-1 mice were injected with LINQ1-tTA 28-d before (A) IVIS imaging of the organs and (B) quantification of luminescence signal. (C) LC-1 mice injected with LINQ1-tTA 24 h post-injection, and (D) organs were excised for (D) assessment of luminescent signal.

